# Computational prediction of cyclotides from *Viola odorata* as potential inhibitors against the neuraminidase of *Streptococcus pneumoniae*

**DOI:** 10.1101/2024.12.08.627407

**Authors:** Sreejanani Sankar, Ajaya Kumar Sahoo, Shanmuga Priya Baskaran, R. Babu, Smita Srivastava, Areejit Samal

## Abstract

Cyclotides are naturally occurring peptides characterized by a cyclic cystine knot, which provides them with exceptional structural stability. In addition to their stability, cyclotides exhibit diverse therapeutic activities including antimicrobial, antiviral and antitumor activities, making them promising candidates in drug discovery. However, computational studies aimed at identifying cyclotide-based inhibitors for infectious diseases remain limited. To this end, we conducted a virtual screening of cyclotides from an Indian medicinal plant *Viola odorata* to identify potential inhibitors against a bacterial pathogen causing respiratory infections. We compiled a library of 93 cyclotides by retrieving their structures from public domain or predicting them using the AlphaFold server. We then docked these cyclotides against the neuraminidase protein of *Streptococcus pneumoniae* and analyzed the interacting residues and binding energies to identify top five potential inhibitors namely, kalata S, kalata B1, cycloviolacin O15, vodo L12, and cycloviolacin O36. Thereafter, we performed molecular dynamics simulations of the protein-cyclotide complexes, and observed that the cyclotides remained stable within the complex. Altogether, this study is the first computational effort to identify potential cyclotide inhibitors against bacterial pathogen causing respiratory diseases, which can be further pursued for experimental validation to develop novel therapeutic agents for respiratory infections.

## Introduction

Cyclotides are a unique class of naturally occurring peptides characterized by a knotted arrangement of three disulfide bonds known as cyclic cystine knot. This unique structural feature of cyclotides provide them with remarkable stability against proteolytic, chemical and thermal degradation [1–3]. In addition to their inherent stability, cyclotides exhibit a range of therapeutic properties including antimicrobial, antiviral and antitumor activities [1, 4–6]. Moreover, cyclotides have a natural role in plant host defense mechanisms [2, 7]. From the perspective of drug discovery, in comparison to the small molecule-based inhibitors, larger surface area of cyclotides offers a greater ability to disrupt interactions between host and pathogen proteins [8, 9]. Importantly, there is growing interest in developing cyclotide-based scaffolds for drug discovery applications due to their efficient synthesis and wide therapeutic potential [10].

To date, CyBase [11] (https://www.cybase.org.au/), the most comprehensive *in silico* database on cyclic proteins, has cataloged over 1300 cyclotides across 90 different plant species, including synthetic constructs. Notably, the plant family Violaceae exhibits the highest abundance of naturally occurring cyclotides [12]. In particular, *Viola odorata* (commonly known as Banafsha), an Indian medicinal plant from the family Violaceae, has long been used to treat ailments such as cold, cough, headache, bronchitis, and fever in traditional systems of medicine [13]. According to the National Medicinal Plants Board, Government of India, the annual trade of *V. odorata* in India is estimated to be 50-100 metric tons [14] (https://nmpb.nic.in/medicinal_list?trade=2334). This demand due to its wide medicinal applications could be attributed to the wide array of bioactive cyclotides produced by *V. odorata*, as confirmed by various experimental studies [13–18]. In spite of the therapeutic potential of plant cyclotides, *in silico* studies to identify potential cyclotide-based inhibitors remain very limited. A notable exception is the computational study aimed at understanding the effect of Cter-M cyclotide from the plant *Clitoria ternatea* on the conformational dynamics of the β-amyloid peptide, which is a characteristic of Alzheimer’s disease [19].

*Streptococcus pneumoniae* is a Gram-positive bacterium that causes pneumonia, bacterial meningitis, respiratory tract infections, septicemia, and otitis media [20, 21]. Notably, India accounts for 20% of childhood mortality worldwide due to pneumonia [22]. Importantly, *S. pneumoniae* is classified as a medium priority drug-resistant bacterium threatening public health, according to the World Health Organization (WHO) Bacterial Priority Pathogens List, 2024 [23, 24]. To this end, peptide-based therapeutics are being explored as alternatives to the conventional small molecule-based antimicrobials [25]. In particular, cyclotide-based inhibitors can be promising candidates in development of antimicrobial drugs with unique mechanisms of action [26]. Notably, previously published studies have shown that extracts of *V. odorata* exhibit antibacterial activity against respiratory pathogens, specifically *S. pneumoniae* [14, 27].

Among the various proteins of *S. pneumoniae*, neuraminidase plays a key role in the bacterial colonization of the mucosal surface in the human upper respiratory tract [28]. *S. pneumoniae* encodes three different neuraminidase proteins namely, NanA, NanB, and NanC, of which NanA is present in all strains and exhibits the highest enzymatic activity [28, 29]. NanA consists of an N-terminal lectin domain, a catalytic domain, and a C-terminal tail which anchors the protein to the bacterial cell wall [29, 30]. The lectin domain recognizes sialic acid from the host cell receptor, while the catalytic domain cleaves the sialic acid, which is further used as nutrient by the bacteria [29, 30]. This process also promotes bacterial growth, adherence and biofilm formation [28–30]. Thus, NanA is a promising target for the development of antimicrobial agents. Though, there have been efforts to adapt influenza virus inhibitors to target NanA protein of *S. pneumoniae* [20], none of the previously published studies have explored cyclotide-based inhibitors against *S. pneumoniae* infection.

Computational approaches can accelerate peptide-based drug discovery by predicting peptide structures, target protein interactions, and binding affinities, thereby reducing the need for extensive experimental screening. Additionally, such methods allow for large-scale screening of peptide libraries to identify promising lead candidates for further validation. In this *in silico* study, we screened a library of cyclotides specific to *V. odorata* to identify potential inhibitors against *S. pneumoniae*. First, we compiled the sequences of cyclotides produced by *V. odorata* as documented in CyBase [11] database, and thereafter, predicted their three-dimensional (3D) structures. Subsequently, we employed computational approaches, including molecular docking and molecular dynamics (MD) simulation, to predict potential cyclotide-based inhibitors against the NanA protein of *S. pneumoniae*. In sum, this is the first computational study undertaken to predict cyclotide-based inhibitors against *S. pneumoniae* infection.

## Methods

### Compilation of ligand library of cyclotides from *Viola odorata*

Cyclotides are a unique class of peptides characterized by cyclic backbones and cyclic cystine knot motifs, which provide them exceptional structural stability [1, 31]. Despite the promising therapeutic properties of cyclotides, including antiviral, anticancer, and antimicrobial activities [1, 32], there have been limited computational efforts [19] to identify cyclotide-based inhibitors targeting specific proteins. Here, we compiled a list of cyclotides documented to be produced by *Viola odorata* (*V. odorata*), an Indian medicinal plant used in traditional systems of medicine [13]. Further, we utilized this list of cyclotides and performed a virtual screening study to identify potential cyclotide inhibitors of a target protein. To obtain the list of cyclotides, we relied on CyBase [11], a comprehensive database dedicated to structurally stable circular peptides, including cyclotides, from various organisms.

First, we collected the amino acid sequences of 96 cyclotides of *V. odorata* from the CyBase database. Then, we were able to obtain the three-dimensional (3D) nuclear magnetic resonance (NMR) structures for 7 of these 96 cyclotides from the RCSB Protein Data Bank (PDB) (https://www.rcsb.org/). For each of the 7 cyclotides, we selected the best NMR model structure based on the lowest Ramachandran distribution Z-score, as predicted by the MolProbity server [33] (http://molprobity.biochem.duke.edu/). For the remaining 89 cyclotides, we predicted the 3D structures from their amino acid sequences using the AlphaFold 3 server [34] (https://alphafoldserver.com/). For each cyclotide sequence, AlphaFold 3 generated five models, and we selected the one with the highest predicted template modelling (pTM) score as the best modelled structure. Following the recommendations by AlphaFold 3 [34], we excluded models with pTM scores below 0.5, leading to the rejection of the predicted structures for 3 cyclotides. In addition, we checked for the presence of three disulfide bonds in the predicted cyclotide structures using an in-house python script. Subsequently, we used the remaining 93 cyclotides, with their structure information, for docking studies (Supplementary Table S1).

### Target protein structure

In this study, we considered neuraminidase protein of *Streptococcus pneumoniae* (*Sp*NanA) as the target protein. Among the many available crystal structures for *Sp*NanA protein in RCSB PDB (https://www.rcsb.org/), we chose the structure with the highest resolution (PDB: 2YA4, 1.8 Å). This structure consists solely the catalytic domain of the neuraminidase protein, which is known to interact with various substrates and small molecule inhibitors [20, 35, 36] (Figure 1). Additionally, the active site residues (R332, R648, R706) are readily accessible within the surrounding environment [20] (Figure 1).

**Figure 1:**
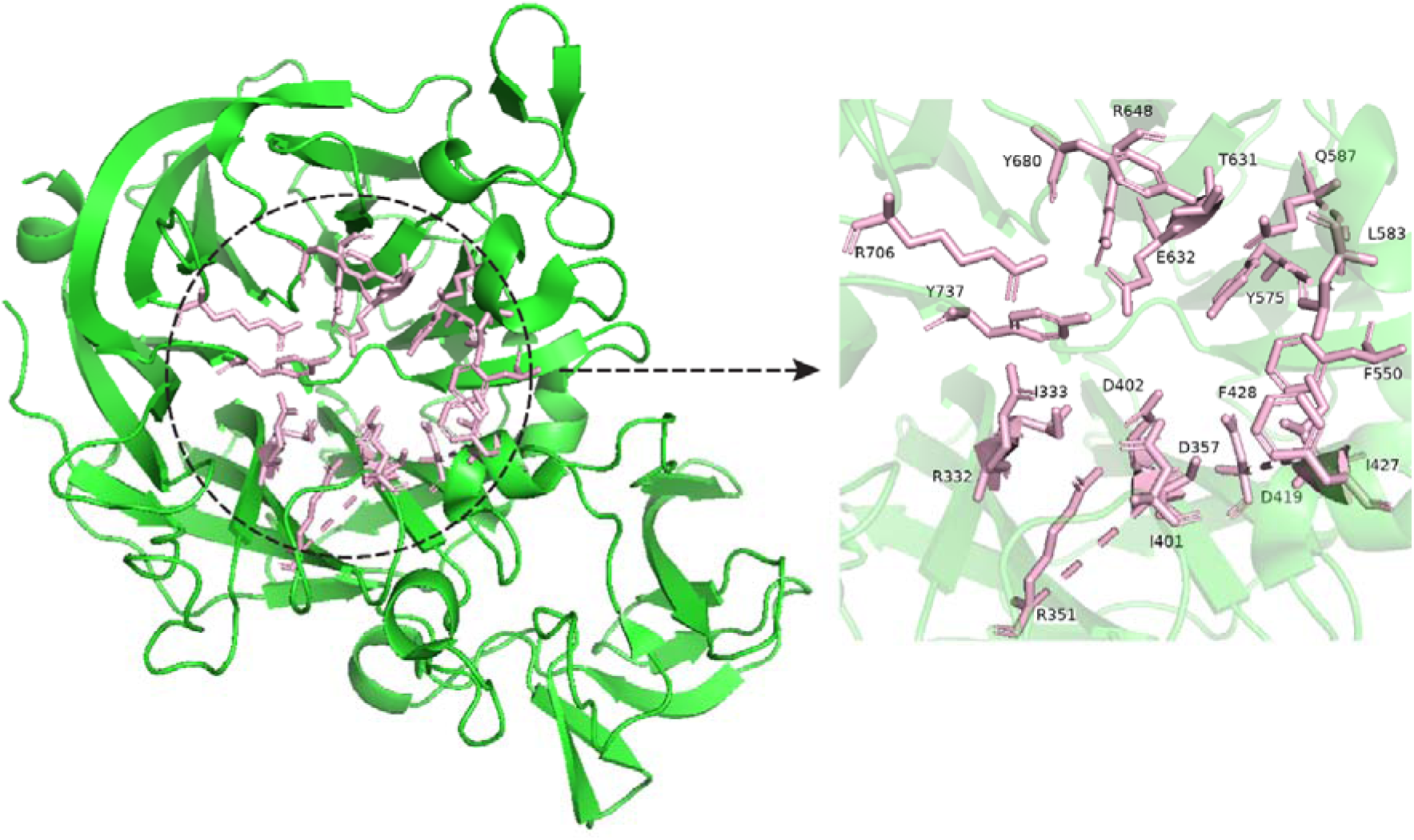
Cartoon representation of the crystal structure of *Sp*NanA. The key amino acid residues of the protein known to interact with small molecule inhibitors namely, R332, I333, R351, D357, I401, D402, D419, I427, F428, F550, Y575, L583, Q587, T631, E632, R648, Y680, R706, and Y737 are shown as pink color sticks. An expanded view showing these key residues is also provided for better visualization.

### Protein-cyclotide docking

To predict potential cyclotide inhibitors of the neuraminidase protein, we performed protein-peptide docking using the FRODOCK 2.0 server [37] (https://frodock.iqf.csic.es/). We docked 93 cyclotides of *V. odorata* against the target protein *Sp*NanA. In the FRODOCK server, the protein structure was uploaded in the ‘Receptor’ panel, cyclotide structure in ‘Ligand’ panel and ‘Others’ was selected in ‘Types of interaction’ panel to perform docking of the cyclotide against the target protein. After completion of the protein-cyclotide docking in the FRODOCK server, we retrieved the ‘receptor.pdb’ and ‘solution.tar’ files. We extracted 10 different poses for each of the cyclotide from the downloaded ‘solution.tar’ file. Using an in-house python script, we combined the receptor and ligand structure files that were retrieved from the FRODOCK server, and thereafter, used the PPCheck (https://caps.ncbs.res.in/ppcheck/) server [38] to identify the best docked pose. In the PPCheck server, we used the ‘Multiple decoys from many different proteins’ option under the ‘Prediction of Right Docking Pose’ panel, to identify the best docked pose among the protein-cyclotide complexes based on their normalized energy per residue value. For the best docked pose, we also used the PPCheck server to compute the strength and type of protein-cyclotide interactions, thereby identifying the binding site residues and their interactions with the cyclotides.

### Molecular dynamics simulation

To assess the stability of the protein-cyclotide complex, molecular dynamics (MD) simulation was performed in the GROMACS version 2024.1 [39, 40]. The AMBER99SB-ILDN force field [41] was used to assign parameters to different amino acids. Both systems were solvated using the SPC216 solvent model and the pH of the system was neutralized by adding necessary Na^+^ and Cl^-^ ions. The structures were placed in a cubic solvent box maintaining a minimum distance of 10 Å between the protein and the box edges. All simulations were performed using the periodic boundary conditions and the electrostatic interactions were calculated using the Particle Mesh Ewald method. The energy of the system was minimized using the steepest descent algorithm for 50000 steps, followed by 100 ps of NVT equilibration at 300 K and NPT equilibration at 1 bar. Throughout these equilibration steps, LINCS [42] algorithm was used to constrain the bond lengths between atoms. Finally protein-ligand complex structure was simulated in triplicate for 500 ns using the Leap-frog algorithm [43], after removing all position restraints. We applied the v-rescale thermostat [44], a modified Berendsen thermostat algorithm for temperature coupling and the Parrinello-Rahman algorithm [45] for pressure coupling during NPT, NVT, and MD simulation.

We utilized GROMACS to compute the root mean square deviation (RMSD), root mean square fluctuation (RMSF), and radius of gyration (R_g_) of the protein-cyclotide complexes. We visualized the MD trajectories in PyMOL [46] software and plotted the graphs using in-house python scripts.

### Free energy computation of protein-cyclotide complex

We computed the binding free energy of the protein-cyclotide complex using the Molecular mechanics/Poisson−Boltzmann surface area (MM-PBSA) method. First, we extracted 201 frames in the interval between 400–500 ns of the MD simulation trajectory for the protein-cyclotide complex. Thereafter, we employed gmx_MMPBSA software [47] to compute the binding free energy of the protein-ligand complexes for the predicted top five cyclotide inhibitors.

## Results and Discussion

### Computational workflow to identify potential cyclotide inhibitors of *Sp*NanA

In this study, we employed a computational workflow to identify potential cyclotide inhibitors of *Sp*NanA (Figure 2). First, we compiled a list of cyclotides of *V. odorata* and their amino acid sequences from the CyBase database. Next, we retrieved their 3D structures from RCSB PDB or predicted their 3D structure using AlphaFold (Methods). Thereafter, we performed protein-peptide docking of 93 cyclotides of *V. odorata* against *Sp*NanA using the FRODOCK web server [37] (Methods). Subsequently, we used the PPCheck web server to identify the best docked pose for each of the cyclotide with *Sp*NanA and computed the interactions between the protein and cyclotide.

**Figure 2:**
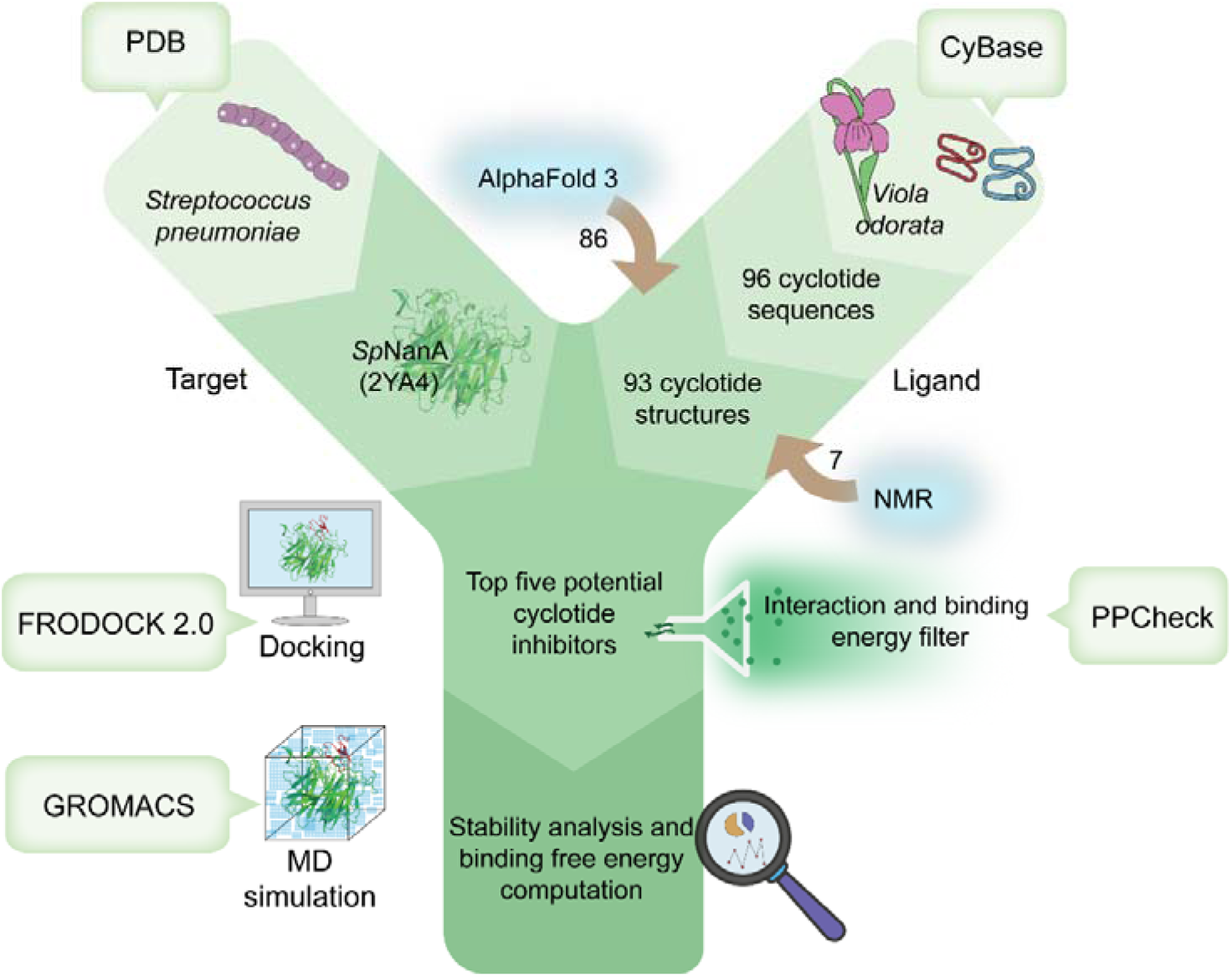
Computational workflow to identify potential cyclotide inhibitors from *Viola odorata* against the neuraminidase protein (*Sp*NanA) of *Streptococcus pneumoniae*.

To shortlist the top cyclotide inhibitors based on the protein-peptide docking, we compared the protein residues interacting with the cyclotide in the best docked pose to a list of 19 residues in *Sp*NanA known to interact with small molecule inhibitors, as reported in published literature [20]. This list of 19 residues in *Sp*NanA include R332, I333, R351, D357, I401, D402, D419, I427, F428, F550, Y575, L583, Q587, T631, E632, R648, Y680, R706, and Y737 (Figure 1). Subsequently, we applied a criterion of at least 12 overlapping residues, i.e., the cyclotide must interact with at least 12 out of the 19 key residues of *Sp*NanA in its best docked pose, to filter cyclotides for further consideration (Supplementary Table S2). Thereafter, we identified the top five cyclotide inhibitors based on the total stabilizing energies (binding energies) of their corresponding protein-ligand complexes as predicted by the PPCheck web server [38] (Supplementary Table S2). Next, we assessed the stability of the protein-cyclotide complex for the top five cyclotides in their best docked pose using MD simulation. Lastly, we computed the free energy of the protein-ligand complex of the cyclotides using gmx_MMPBSA software.

### Prediction of top five cyclotide inhibitors of *Sp*NanA

Using the computational workflow depicted in Figure 2, we predicted top five cyclotide inhibitors of *Sp*NanA namely, kalata S, kalata B1, cycloviolacin O15, vodo L12, and cycloviolacin O36. These inhibitors were selected based on their interactions with the key amino acid residues in the protein structure and their binding energy with the protein, as determined through docking analysis.

The cyclotide C1 (kalata S) has a docking based binding energy of -266.15 kJ/mol (Table 1). Kalata S is known to exhibit immunomodulatory [12] and anti-cancer properties [48]. Among the key amino acid residues in *Sp*NanA known to interact with small molecule inhibitors, R332, L583, R648, Y680 and R706 form short contacts with kalata S (Table 1; Figure 3). Further, R332, I333, R351, D357, I401, D402, I427, F428, Y575, L583, Q587, T631, E632, R648, Y680, R706 and Y737 form van der Waals interactions with the cyclotide (Figure 3).

**Figure 3:**
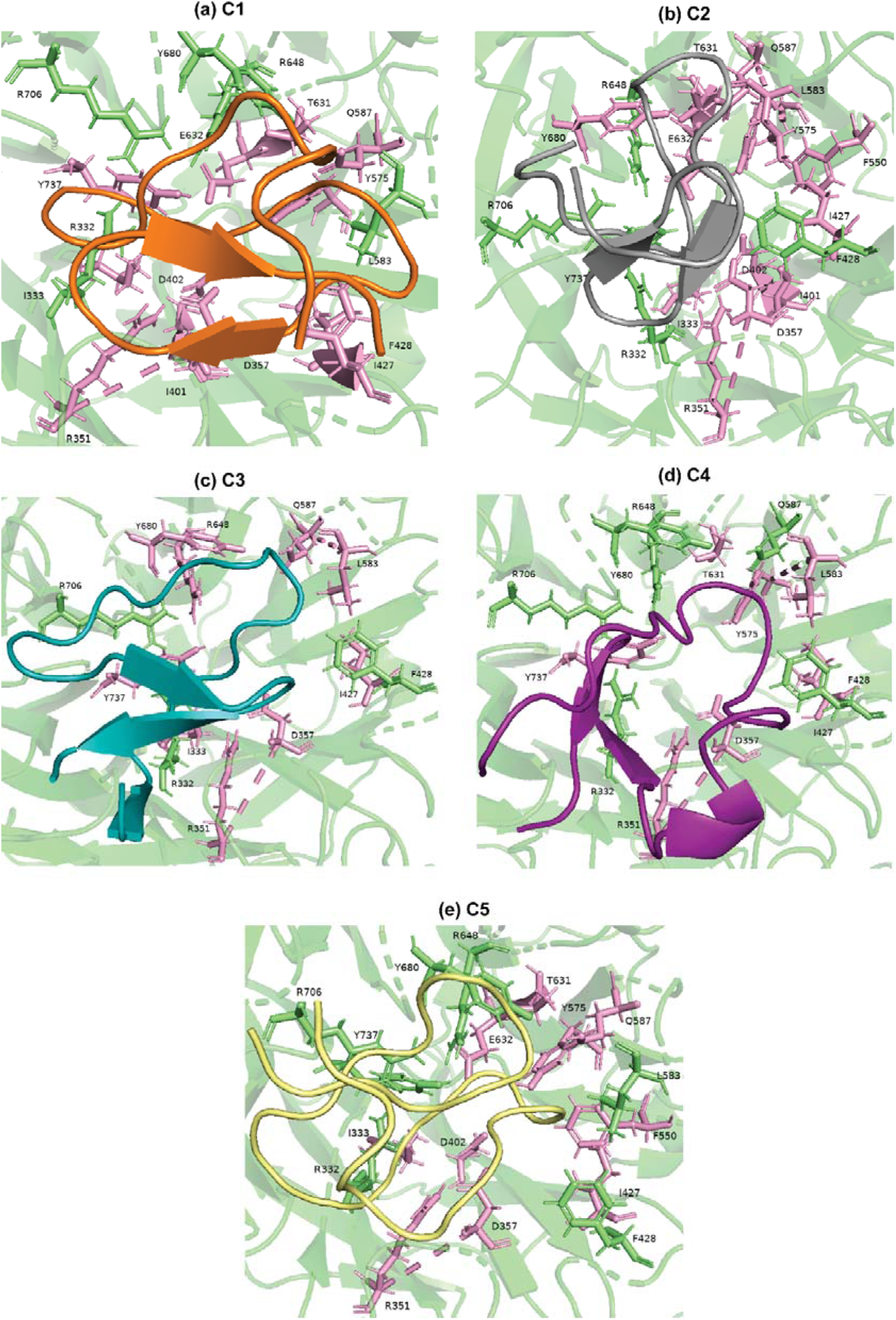
Cartoon representation of non-covalent interaction of the protein *Sp*NanA with the top five predicted cyclotide inhibitors (C1-C5) in their best docked pose. In each case, the amino acid residues shown as pink sticks represent van der Waals interactions with the cyclotide, and the amino acid residues shown as green sticks represent mixed non-covalent interactions with the cyclotide. The atoms in the cyclotides are shown in orange for C1, in grey for C2, blue for C3, magenta for C4, and yellow for C5.

**Table 1:**
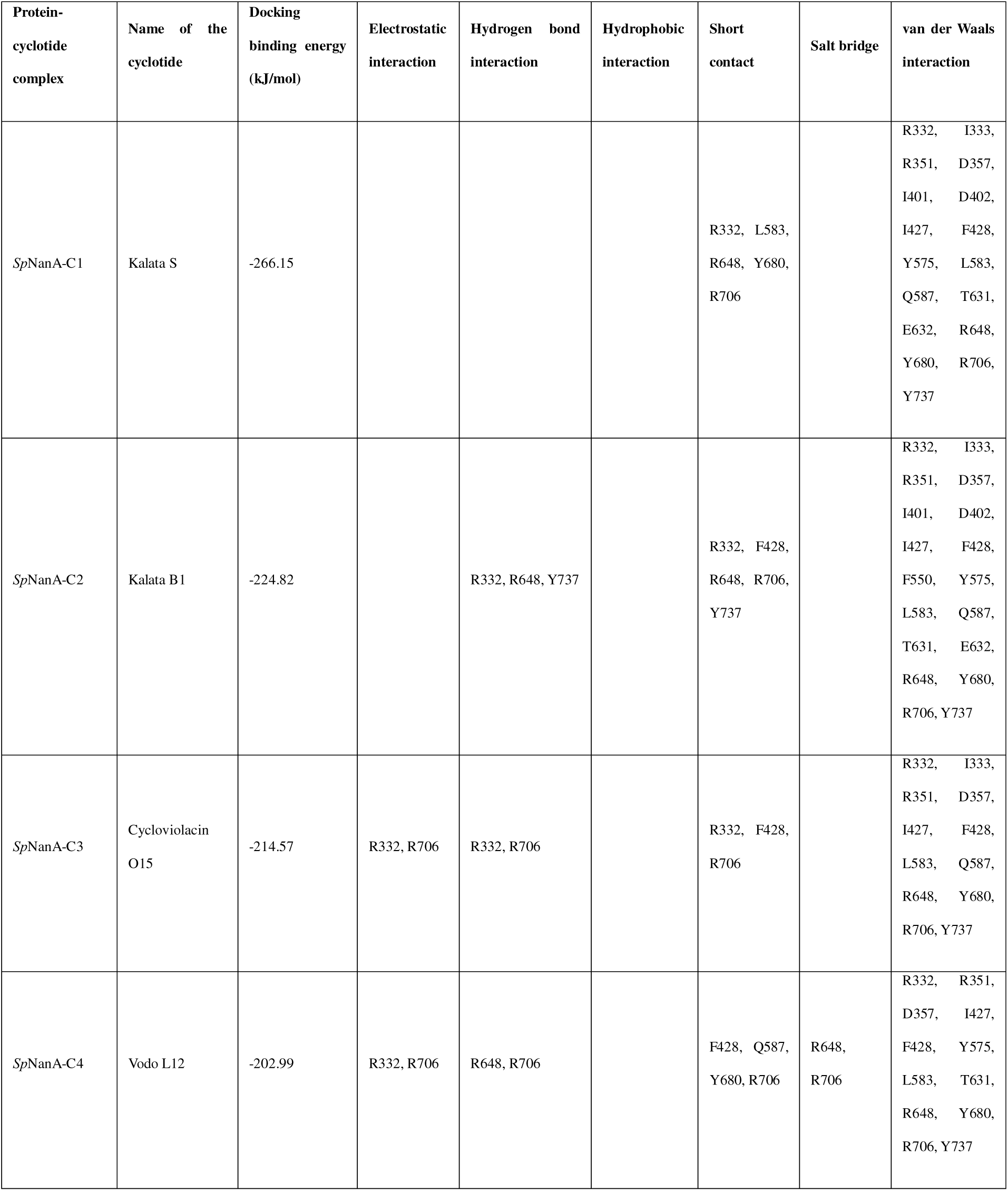
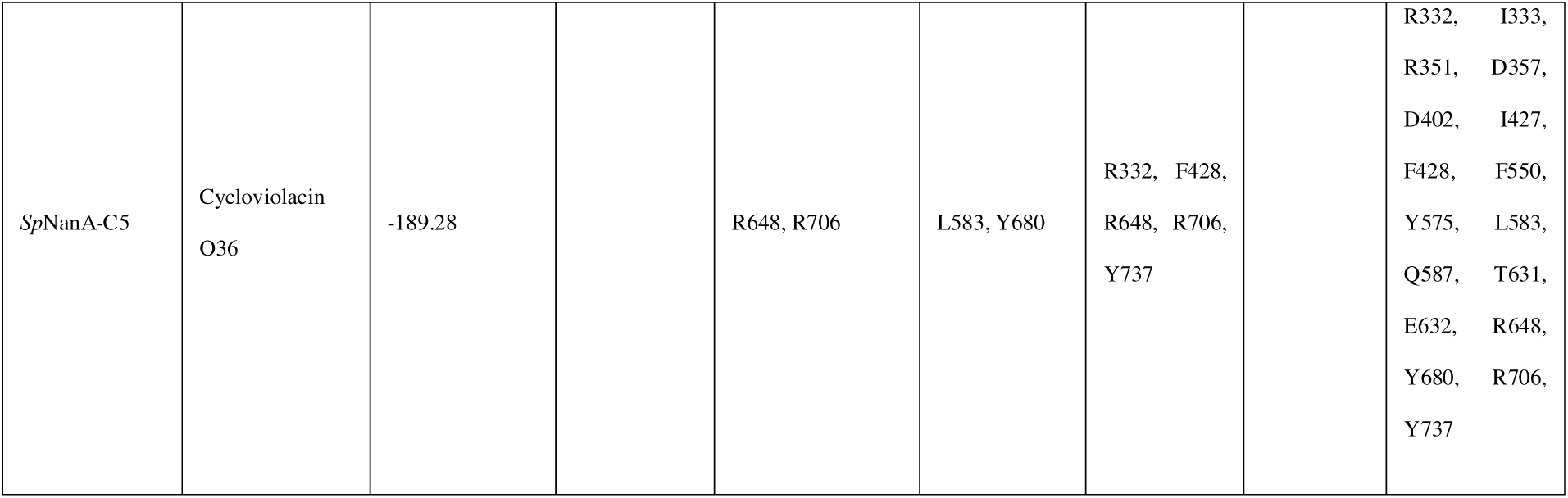
Non-covalent interaction between the amino acid residues of the protein *Sp*NanA and the top five predicted cyclotide inhibitors in the best docked pose, predicted from PPCheck web server.

The cyclotide C2 (kalata B1) has a docking based binding energy of -224.82 kJ/mol (Table 1). Kalata B1 is documented for diverse therapeutic potential including antibacterial, nematicidal, molluscicidal, insecticidal and anti-cancer properties [49–54]. Notably, the plant biomass of *V. odorata* containing higher abundance of kalata B1 has been shown earlier by some of us to exhibit antibacterial activity against *S. pneumoniae* [13, 14]. Among the key amino acid residues in *Sp*NanA known to interact with small molecule inhibitors, R332, R648 and Y737 forms hydrogen bond interactions with kalata B1 (Table 1; Figure 3). Additionally, R332, F428, R648, R706 and Y737 form short contacts with the cyclotide. Further, R332, I333, R351, D357, I401, D402, I427, F428, F550, Y575, L583, Q587, T631, E632, R648, Y680, R706 and Y737 form van der Waals interactions with kalata B1 (Table 1; Figure 3).

The cyclotide C3 (cycloviolacin O15) has a docking based binding energy of -214.57 kJ/mol (Table 1). Cycloviolacin O15 is known to exhibit anthelmintic activity against nematodes [55]. Among the key amino acid residues in *Sp*NanA known to interact with small molecule inhibitors, R332 and R706 form hydrogen bond interactions with cycloviolacin O15 (Table 1; Figure 3). Additionally, R332 and R706 form favorable electrostatic interactions with the cyclotide. R332, F428 and R706 form short contacts with the cyclotide. Further, R332, I333, R351, D357, I427, F428, L583, Q587, R648, Y680, R706 and Y737 form van der Waals interactions with cycloviolacin O15 (Table 1; Figure 3).

The cyclotide C4 (vodo L12) has a docking based binding energy of -202.99 kJ/mol (Table 1). Among the key amino acid residues in *Sp*NanA known to interact with small molecule inhibitors, R648 and R706 form hydrogen bond interactions with the cyclotide. Additionally, R648 and R706 form salt bridges with the cyclotide (Table 1; Figure 3). Both R332 and R706 form electrostatic interactions with Vodo L12. Moreover, F428, Q587, Y680 and R706 form short contacts with the cyclotide (Table 1; Figure 3). Further, R332, R351, D357, I427, F428, Y575, L583, T631, R648, Y680, R706 and Y737 form van der Waals interactions with vodo L12 (Table 1; Figure 3).

The cyclotide C5 (cycloviolacin O36) has a docking based binding energy of -189.28 kJ/mol (Table 1). Among the key amino acid residues in *Sp*NanA known to interact with small molecule inhibitors, R648 and R706 form hydrogen bond interactions with the cyclotide (Table 1; Figure 3). L583 and Y680 form hydrophobic interactions with the cyclotide. R706 form favorable electrostatic interaction with the cyclotide. Moreover, R332, F428, R648, R706 and Y737 form short contacts with the cyclotide (Table 1; Figure 3). Further, R332, I333, R351, D357, D402, I427, F428, F550, Y575, L583, Q587, T631, E632, R648, Y680, R706 and Y737 form van der Waals interactions with the cycloviolacin O36 (Table 1; Figure 3).

### MD based stability analysis of *Sp*NanA in complex with top five potential cyclotide inhibitors

We employed MD simulations to evaluate the stability of the protein-cyclotide complexes for the predicted top five cyclotide inhibitors namely, kalata S (C1), kalata B1 (C2), cycloviolacin O15 (C3), vodo L12 (C4), and cycloviolacin O36 (C5). Each protein-cyclotide complex was subjected to 500 ns MD simulations in duplicate (Methods). The results indicated that all the five cyclotides remained stable in the respective protein-cyclotide complexes (Figure 4).

**Figure 4:**
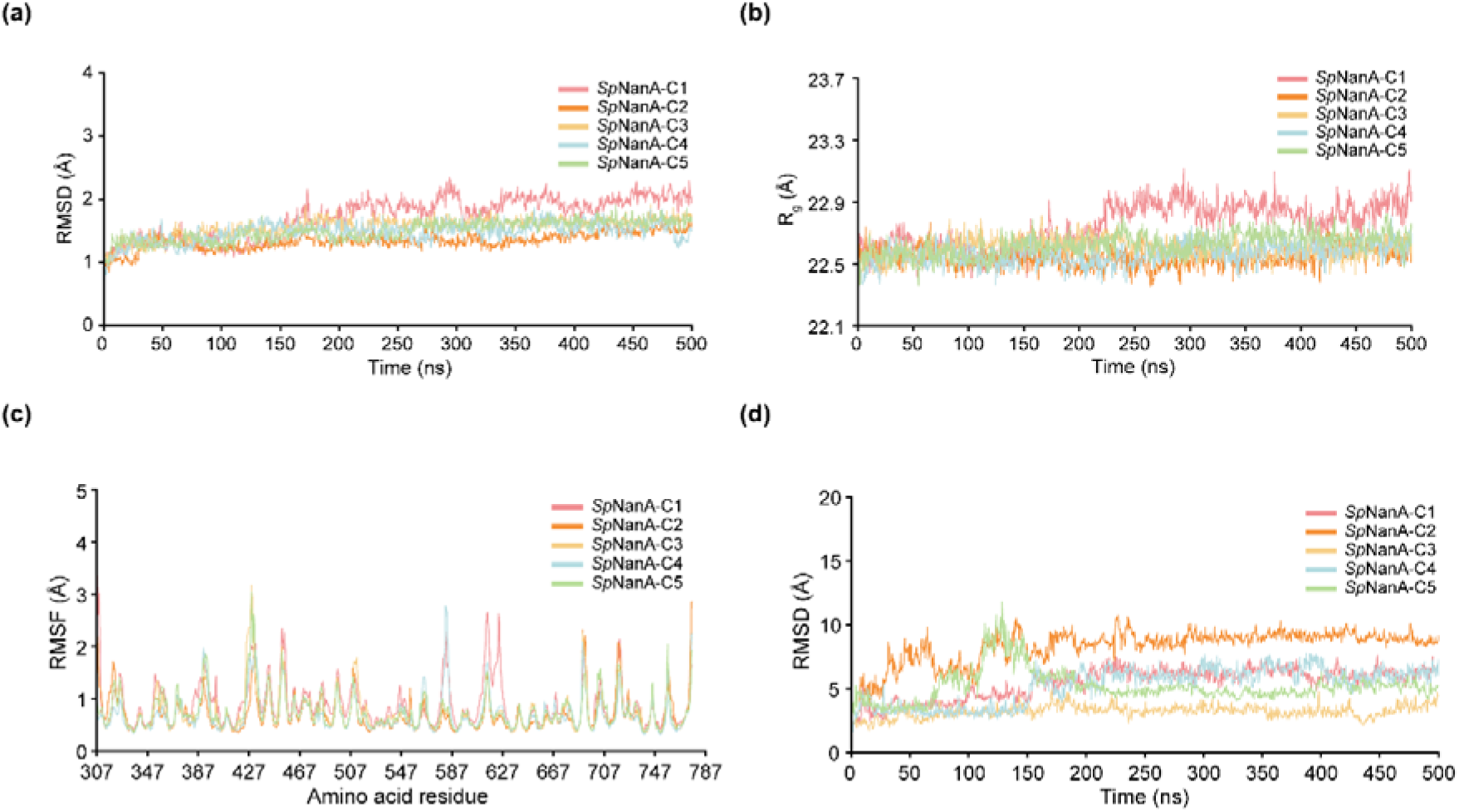
Analysis of MD trajectories from 500 ns simulation of the protein-ligand complexes for the top five predicted cyclotide inhibitors (C1-C5). **(a)** RMSD of the Cα atoms of amino acid residues in *Sp*NanA within the protein-cyclotide complex. **(b)** R_g_ of the *Sp*NanA protein structure within the protein-cyclotide complex. **(c)** RMSF of the Cα atoms of amino acid residues in *Sp*NanA within the protein-cyclotide complex. **(d)** RMSD of the heavy (non-hydrogen) atoms of the cyclotide within the protein-cyclotide complex.

The average RMSD of the Cα atoms of the protein were 1.72 ± 0.32 Å for the *Sp*NanA-C1 complex, 1.34 ± 0.14 Å for the *Sp*NanA-C2 complex, 1.56 ± 0.16 Å for the *Sp*NanA-C3 complex, 1.48 ± 0.16 Å for the *Sp*NanA-C4 complex, and 1.50 ± 0.16 Å for the SpNanA-C5 complex (Figure 4a). Additionally, the R_g_ of the complete protein structure remained stable throughout the simulations (Figure 4b). The average R_g_ values for the protein were 22.75 ± 0.14 Å for the *Sp*NanA-C1 complex, 22.54 ± 0.06 Å for the *Sp*NanA-C2 complex, 22.60 ± 0.06 Å for the *Sp*NanA-C3 complex, 22.62 ± 0.08 Å for the *Sp*NanA-C4 complex, and 22.57 ± 0.07 Å for the *Sp*NanA-C5 complex (Figure 4b). Further, the RMSF of the amino acids of the protein in the protein-cyclotide complexes for the top five cyclotides also showed stability (Figure 4c). Similarly, the RMSD of the heavy atoms for all five cyclotides in the corresponding protein-cyclotide complexes show stability throughout the MD simulation (Figure 4d).

MM-PBSA is a widely used method to accurately estimate the binding free energy of protein-ligand complex [56–58]. We utilized MM-PBSA based approach to compute the binding free energy of the protein-cyclotide complexes for the top five potential cyclotide inhibitors (Methods). The binding free energy were computed to be -16.03 ± 4.82 kcal/mol for the *Sp*NanA-C1 complex, -29.62 ± 4.14 kcal/mol for the *Sp*NanA-C2 complex, -19.20 ± 4.12 kcal/mol for the *Sp*NanA-C3 complex, -44.10 ± 7.02 kcal/mol for the *Sp*NanA-C4 complex, and -15.90 ± 4.53 kcal/mol for the *Sp*NanA-C5 complex.

## Conclusion

Cyclotides are cyclic peptides found in plants, characterized by disulfide bonds that confer exceptional stability, making them promising candidates for drug discovery. However, there has been limited research using computational approaches to identify potential cyclotide inhibitors to combat infectious diseases. To address this, we curated a list of cyclotides known to be produced by an Indian medicinal plant, *V. odorata*, and thereafter, performed a virtual screening to identify potential cyclotide inhibitors against *S. pneumoniae*. We performed molecular docking of 93 cyclotides against the neuraminidase protein of *S. pneumoniae*, and based on their interacting residues and binding energies, identified five potential inhibitors. Further, we assessed the stability of the protein-cyclotide complexes for the top five cyclotide inhibitors and found that all the five cyclotides are stable in the corresponding protein-ligand complexes. To reiterate, this is the first study, which performs large-scale computational screening of cyclotides to identify potential inhibitors against bacterial infection.

Peptide-based inhibitors are emerging as promising alternatives to small molecule inhibitors due to their higher specificity toward target proteins [59]. Their larger size compared to small molecule inhibitors enables them to effectively target and inhibit protein-protein interactions involving larger surface areas [59]. Among these, cyclotides are particularly notable for their exceptional structural stability, making them attractive candidates for therapeutic applications. In this computational study, five cyclotides from the *V. odorata* plant namely, kalata S, kalata B1, cycloviolacin O15, vodo L12, and cycloviolacin O36, were identified as potential inhibitors against *S. pneumoniae*. Supporting this, some of us [14] previously demonstrated that cyclotide rich plant biomass from *V. odorata* is effective against bacteria causing respiratory infections. Altogether, these findings highlight the potential of cyclotides as therapeutic agents, though further cyclotide-specific experimental studies are required to validate the predicted candidates in this computational study as effective inhibitors against respiratory infections.

## Author Contributions

**Sreejanani Sankar:** Conceptualization, Data Curation, Formal Analysis, Methodology, Software, Visualization, Writing; **Ajaya Kumar Sahoo:** Conceptualization, Data Curation, Formal Analysis, Methodology, Software, Visualization, Writing; **Shanmuga Priya Baskaran:** Conceptualization, Data Curation, Formal Analysis, Visualization, Writing; **R. Babu:** Conceptualization, Data Curation, Formal Analysis, Writing; **Smita Srivastava:** Conceptualization, Formal Analysis, Methodology, Supervision, Writing; **Areejit Samal:** Conceptualization, Formal Analysis, Methodology, Supervision, Writing.

## Acknowledgements

Areejit Samal would like to acknowledge funding from the Department of Atomic Energy (DAE), Government of India via Apex Project to The Institute of Mathematical Sciences (IMSc), Chennai. R. Babu would like to acknowledge the Science and Engineering Research Board (SERB), Department of Science and Technology (DST), Government of India, Confederation of Indian Industry (CII) and Himalaya Wellness Company for the award of Prime Minister’s Fellowship for Doctoral Research (SERB/PM Fellow/CII-FICCI/Meeting/2019).

## Declarations

### Conflict of interest

The authors declare that they have no conflicts of interest.

### Ethical approval

This is a purely computational study and ethics approvals are not applicable.

### Consent to participate

Not applicable.

## Consent for publication

All co-authors have read and approved the manuscript.

## Supplementary Table Captions

**Supplementary Table S1:** The table provides a curated list of 93 cyclotides used for computational screening against the neuraminidase protein of *Streptococcus pneumoniae* (*Sp*NanA) to identify potential inhibitors. For each cyclotide, the table provides its name, amino acid sequence, the source of its corresponding three-dimensional (3D) structure (obtained from RCSB PDB or predicted by AlphaFold 3), and the predicted template modelling (pTM) score of the predicted structure.

**Supplementary Table S2:** The table lists the cyclotides that form non-covalent interactions with 12 or more key amino acid residues in the protein *Sp*NanA. For each cyclotide, the table provides the number of key residues in the protein involved in non-covalent interactions with the cyclotide and the binding energy of the protein-cyclotide complex (in kJ/mol) obtained using the PPCheck web server.

## Notes

### Competing Interest Statement

The authors have declared no competing interest.

